# Simultaneous application of enzyme and thermodynamic constraints to metabolic models using an updated Python implementation of GECKO

**DOI:** 10.1101/2023.03.20.533446

**Authors:** Jorge Carrasco Muriel, Christopher Long, Nikolaus Sonnenschein

## Abstract

GEnome-scale Metabolic (GEM) models are knowledge bases of the reactions and metabolites of a particular organism. These GEM models allow for the simulation of the metabolism - e.g. calculating growth and production yields - based on the stoichiometry, reaction directionality and uptake rates of the metabolic network. Over the years, several extensions have been added to take into account other actors in metabolism, going beyond pure stoichiometry. One such extension is enzyme-constraint models, which enable the integration of kinetic data and proteomics data into GEM models. Given its relatively recent formulation, there are still challenges in standardization and data reconciliation between the model and the experimental measurements. In this work, we present geckopy 3.0 (Genome-scale model Enzyme Constraints, using Kinetics and Omics in python), an actualization from scratch of the previous python implementation of the same name. This update tackles the aforementioned challenges, in an effort to reach maturity in enzyme-constraint modeling. With the new geckopy, proteins are typed in the SBML document, taking advantage of the SBML Groups extension, in compliance with community standards. Additionally, a suite of relaxation algorithms - in the form of linear and mixed-integer linear programming problems - has been added to facilitate reconciliation of raw proteomics data with the metabolic model. Several functionalities to integrate experimental data were implemented, including an interface layer with pytfa for the usage of thermodynamics and metabolomics constraints. Finally, the relaxation algorithms were benchmarked against public proteomics datasets in *Escherichia coli* for different conditions, revealing targets for improving the enzyme constrained model and/or the proteomics pipeline.

**IMPORTANCE:** The metabolism of biological cells is an intricate network of reactions that interconvert chemical compounds, gathering energy and using that energy to grow. The static analysis of these metabolic networks can be turned into a computational model which is able to efficiently output the distribution of fluxes in the network. With the inclusion of enzymes in the network, we can also interpret the role and concentrations of the metabolic proteins. However, the models and the experimental data often clash, resulting in a network that cannot grow. Here, we tackle this situation with a suite of relaxations algorithms in a package called geckopy. Additionally, to ensure that enzyme-constraint models follow the community standards, a format for the proteins is postulated. Geckopy also integrates with other software to allow for adding thermodynamic and metabolomic constraints. We hope that the package and algorithms presented here will serve useful for the constraint-based modeling community.

## Introduction

For those who have taken courses in Biochemistry, Metabolism and metabolic regulation as undergraduates, the calculation of yields of ATP from a substrate may sound quite familiar. They may remember the effort of memorizing the balance of the different routes in the central metabolism, how these metabolic routes branch out and the difficulty of clarifying the destiny of the flux of carbon, as it is distributed across the network.

The complexity and ubiquity of these calculations in Biochemistry curricula was noted by M. R. Watson in the 80s [1]. He devised how this process, which he regarded as laborious, could be handled by a computer, both easing the understanding of Metabolism and demonstrating the application of computation in Biochemistry.

Thus, reactions can be rewritten as computations. Consider, for instance Eq 1. In this equation, we have two reactions and 4 metabolites {*A, B, C, D*}.

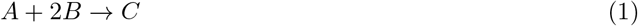

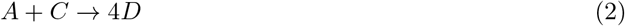

We can rewrite the reactions as a matrix **S** using the stoichiometric coefficients Eq 3.

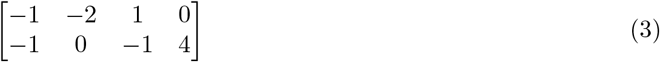

Now, by enforcing mass conservation over the network, **Sv** = 0, we can convert the metabolic network into a linear programming problem [2]. By optimizing the so-called biomass function [3], i.e., an artificial sink reaction added to the metabolism network that turns essential precursor metabolites into biomass based on empirically determined proportions, growth can be simulated. When the reactions and metabolites of a whole species are gathered into one of these models, we call it a GEnome-scale Metabolic (GEM) model. GEMs are stored in the Systems Biology Markup Language (SBML) [4], which is an open standard for the Systems Biology community.

Far from being mere learning tools in the classrooms, Genome-scale Metabolic Models have proven to be useful in a variety of ways, from strain-design for biochemical production, drug-target prediction [5] and improving the understanding of human diseases and the metabolism of different organisms [6].

In addition, GEM models can be used as a platform to integrate omics data. GEM models can be extended to include turnover numbers *k_cat_* and enzyme concentrations [7,8] by including proteins as pseudometabolites in the reactions (Eq 4).

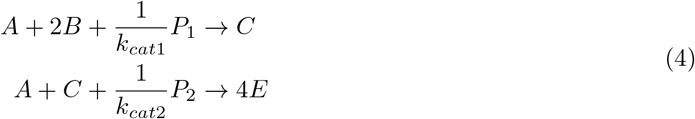

Hence, enzyme-constraint models expand the stoichiometric matrix **S** by adding new protein ”metabolites” *P*. Furthermore, exchange pseudorreactions are added to supply the proteins into the network. The supply is constrained to an upper bound, which may be used to indicate an experimental measurement of the given protein.

Additionally, a formulation to include thermodynamics called Thermodynamics Flux Analysis (TFA) has also been proposed and implemented in previous publications [9,10]. With the inclusion of reaction Gibbs free energies, TFA constrains the reactions to follow the second law of thermodynamics. Conveniently, metabolomics concentrations can be integrated in the computation of Gibbs free energies.

Typing of proteins in the SBML document remains a challenge. Until now, these enzymes have been stored as *SBML Species*, relying on naming conventions to differentiate them from metabolites. This is error-prone and also hinders their usage within existing COnstraints-Based Reconstruction and Analysis (COBRA) software [11–13]. Moreover, the integration of absolute proteomics in enzyme-constraint models is not trivial; it usually yields models that are infeasible for the given reported growth and exchange fluxes, requiring some relaxation of the experimental constraints [14].

In this work, we present an update of an existing open-source enzyme-constraint software: geckopy 3.0. This update tackles the aforementioned problems, providing 1) a SBML-compliant formulation of proteins; 2) relaxation algorithms for reconciling the model with the experimental data; and 3) an integration layer with existing academic software - pytfa [10] - to include thermodynamic and metabolomics constraints [9] on top of enzyme-constraint models. Finally, we benchmark the assumptions and performance of the relaxation algorithms here presented and inspect the effects of the different layered constraints over the plain stoichiometric formulation, highlighting the difficulty and caveats of integrating different types of omics in the same model.

## Materials and methods

Two jupyter notebooks reproducing the results in this paper are provided as supplementary material: S2 File corresponds to Figures 1, 2, 3, 4; and S3 File corresponds to Figure 5.

**Fig 1.**
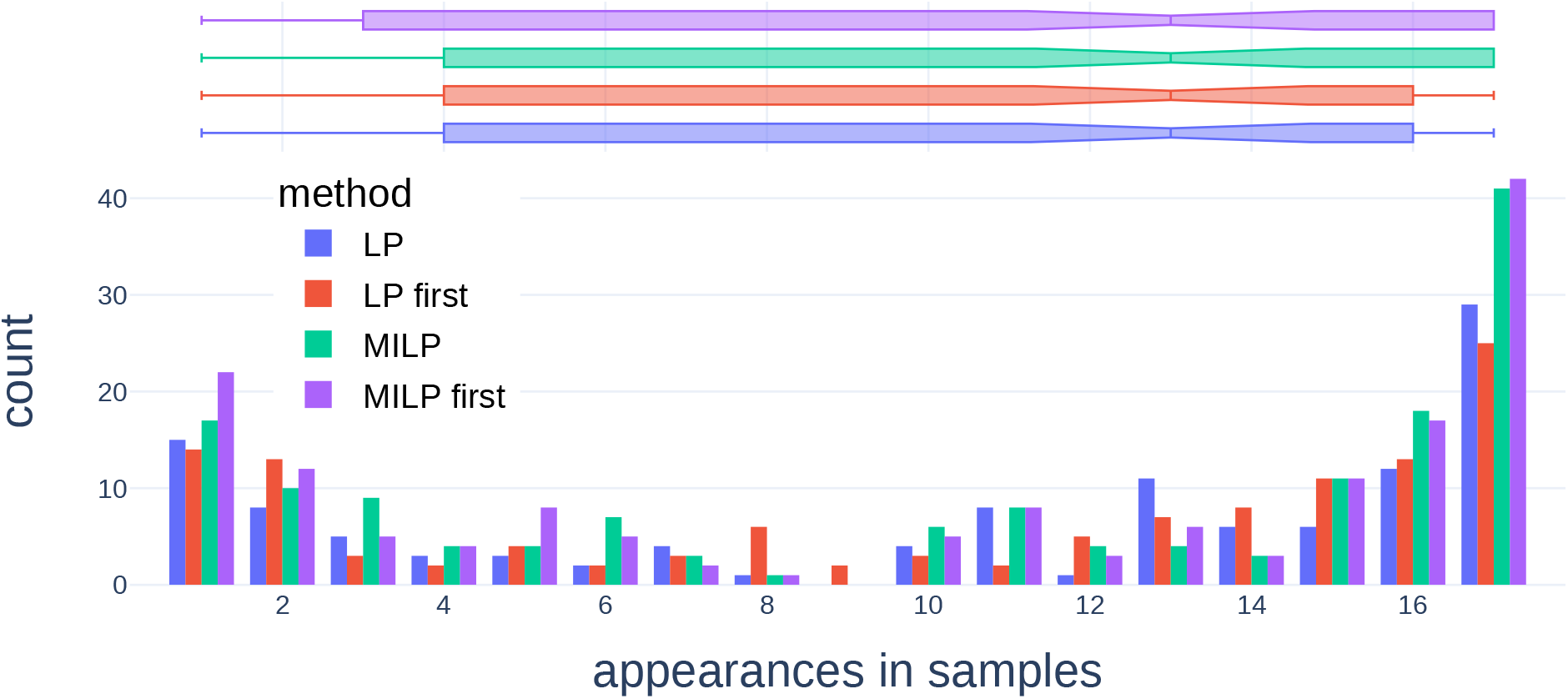
Number of samples where a protein was included in the IIS for the different relaxation algorithms, over 17 samples. The ”first” alternatives correspond to running the relaxation just once without expanding the IIS.

**Fig 2.**
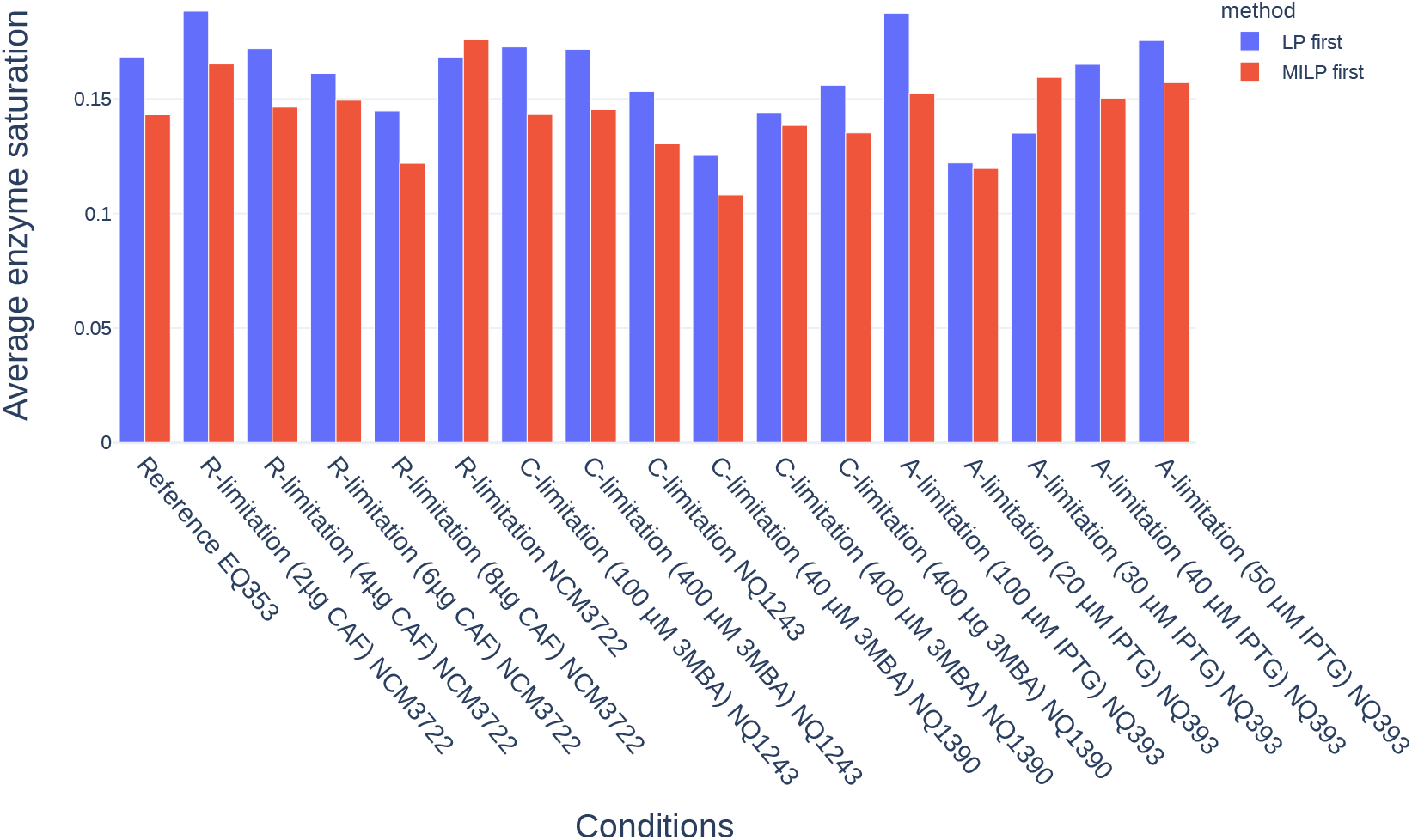
Average enzyme saturation in terms of the protein usage (flux) of each protein divided by its concentration per condition.

**Fig 3.**
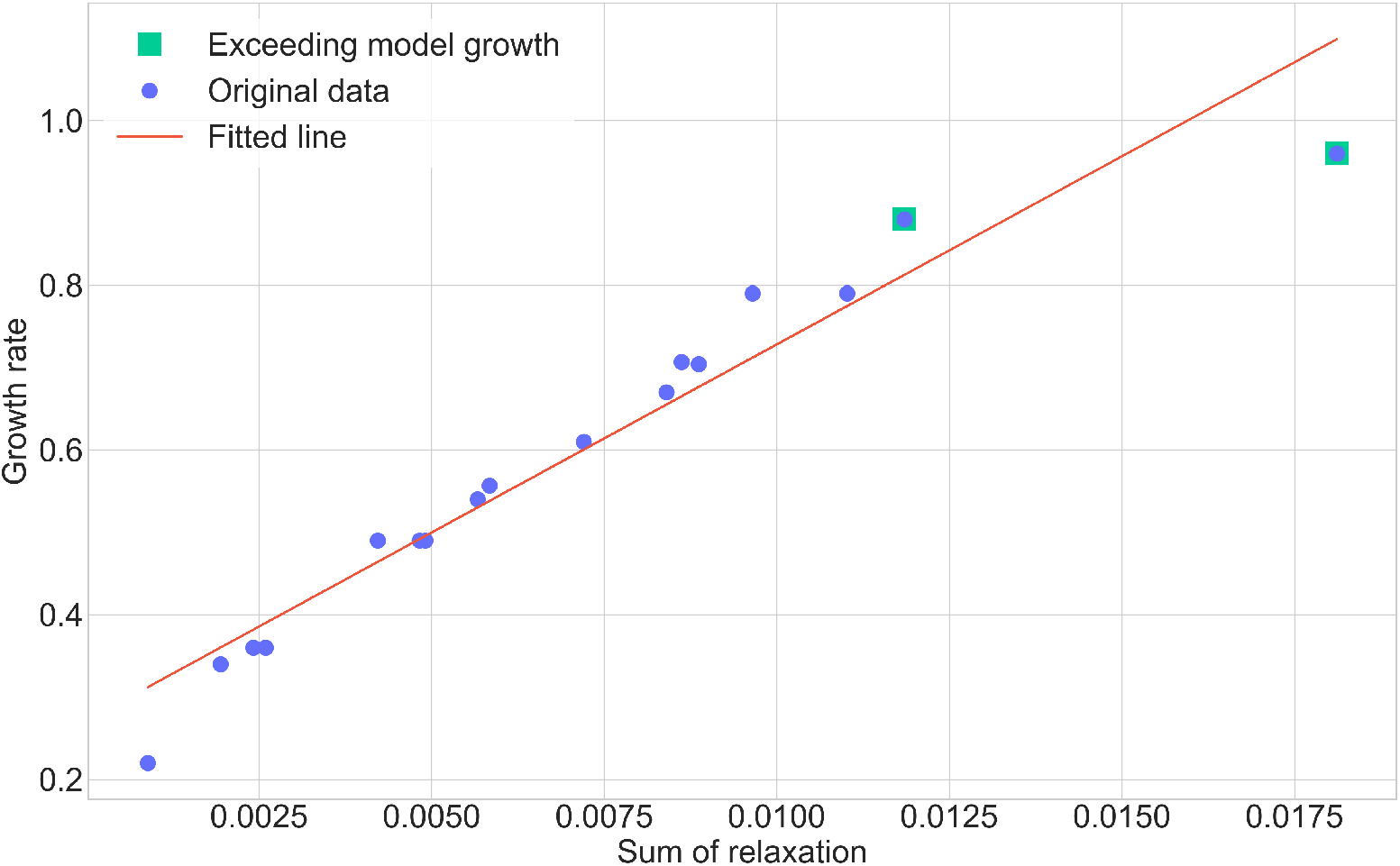
Least squares fitting from the sum of the relaxation values for each proteomics condition with respect to the growth rate. The Pearson’s correlation coefficient was 0.966. The squares represent the samples that exceeded the growth rate of and that were capped to the unconstrained growth of the model. The rightest condition, which deviates from the fitting, is the R-limited positive control.

**Fig 4.**
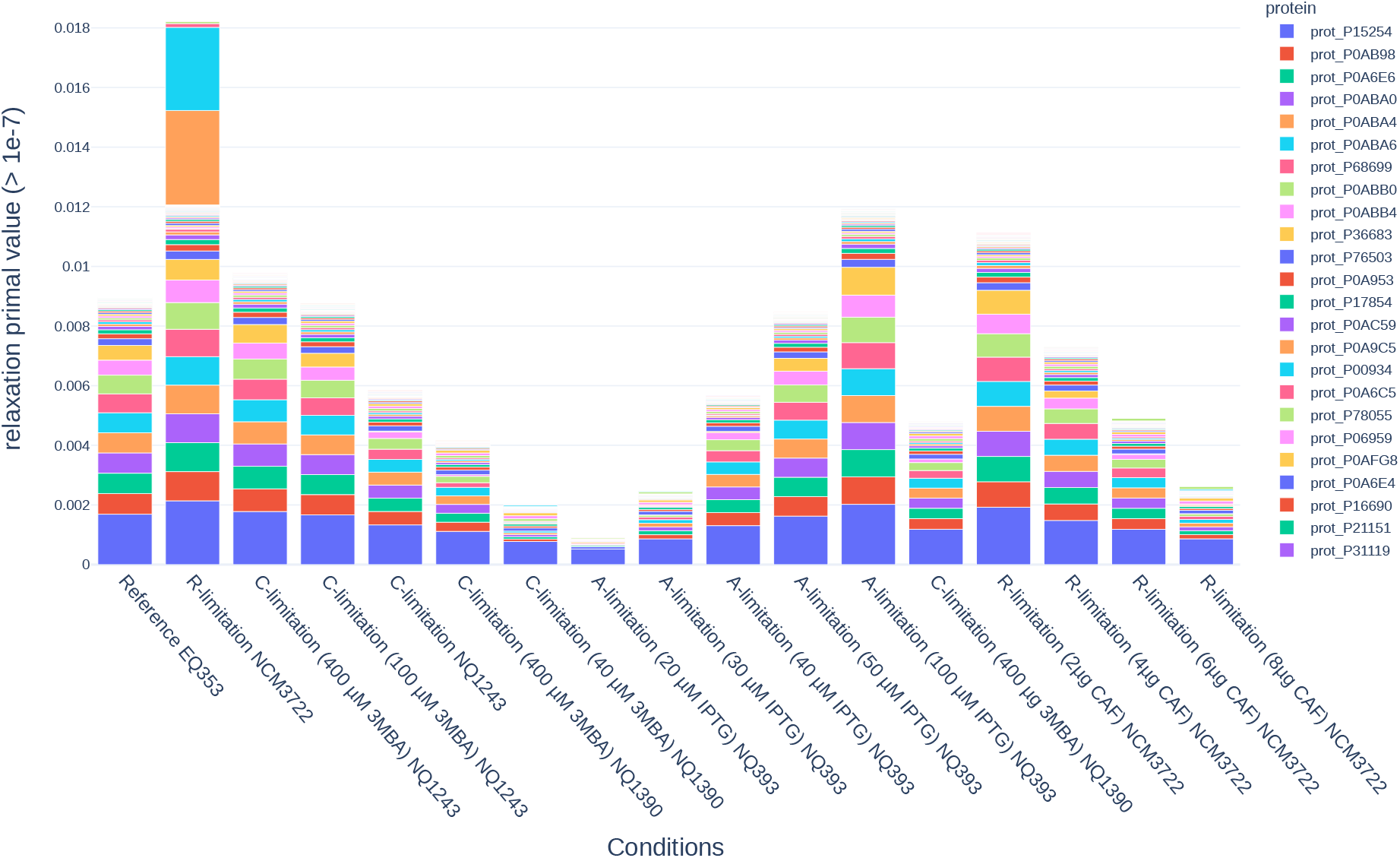
Protein relaxation value (as primal values, in *mmol/g_DW_/h*) for the proteins of the IIS solved by the LP with one iteration per condition (LP first). Only relaxation values greater than 10^-7^ (in the numerical error threshold for the LP solvers) were reported.

**Fig 5.**
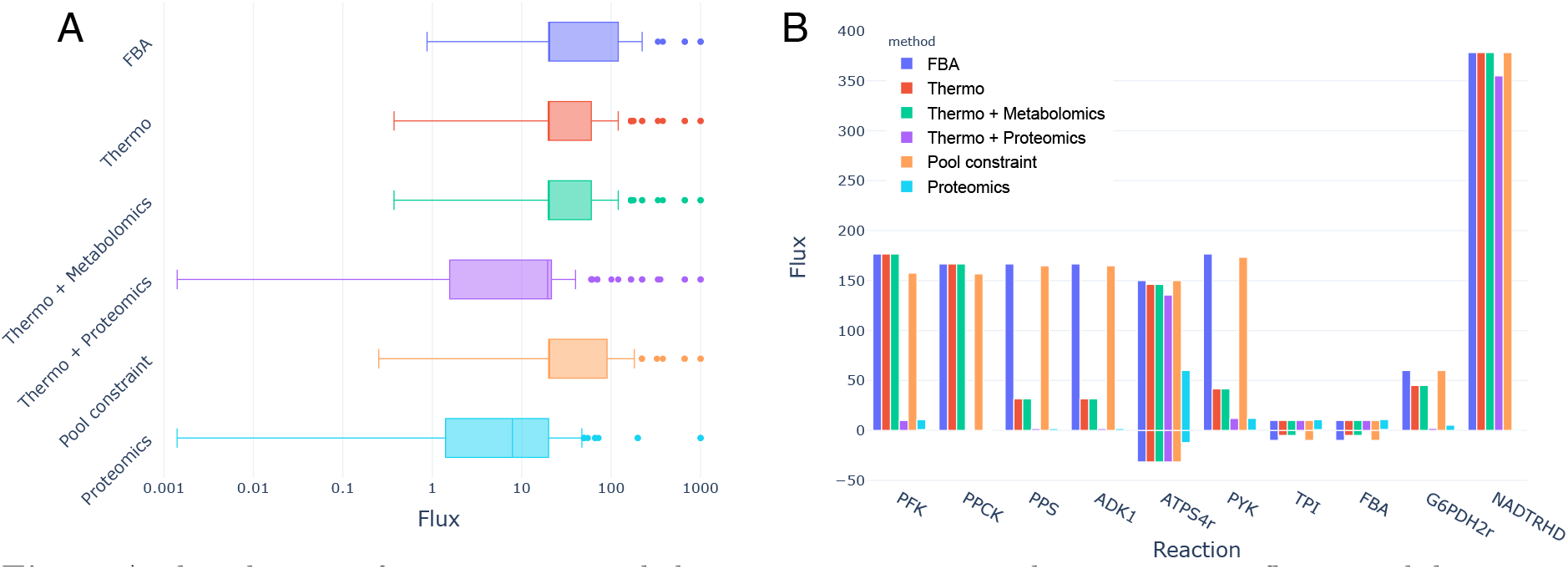
A: distribution of non-zero intervals between maximums and minimums in flux variability analysis of the *E. coli* core model, in logarithmic scale. Overall, protein constraints produce tighter distributions, with lower values of the interval. B: FVA maxima and minima for some of the representative reactions in the model.

### Flux balance analysis

Flux Balance Analysis (FBA) is a linear programming problem in the form of equation Eq 5 [2].

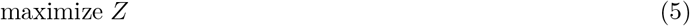

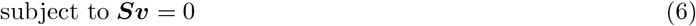

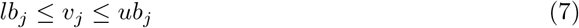

where ***Z*** is the objective, ***S*** is the stoichiometric matrix; ***v*** is the vector of reaction variables; and *lb_j_* and *ub_j_* are the lower and upper bounds constraining the flux of the reaction variable *v_j_*.

***Z*** can be expressed as ***c**^T^**v*** where ***c*** is the coefficient vector that indicates the influence of each reaction in ***v*** over the objective. Typically, ***c*** is a standard basis vector referring to only one reaction, the biomass pseudorreaction or the ATP synthase. The biomass pseudorreaction acts as a sink in the metabolism, converting different byproducts of the metabolic network into mass per time (generally *g/g_DW_/h*), calibrated by experimental data.

Eq 6 constitutes the steady-state assumption, which forces a net balance of fluxes through the network. This produces an undetermined system of equations with infinite solutions (since there are more reactions than metabolites), whose solution space is further constrained by Eq 7. The latter equation imposes the minimum and maximum flux value for each reaction in *V*. In big GEM models, the solution of fluxes that optimizes 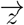 is generally non-unique.

### Enzyme-constraint Flux balance analysis

Enzyme-constraint Flux Balance Analysis is another linear programming problem built on top of FBA. For a more detailed explanation, refer to [7,8]. Briefly, the stoichiometric matrix is extended with protein exchanges (variables/columns) and protein species *P* (constraints/rows).

In the enzyme-constraint formulation, a given enzyme *p* participates in its respective reaction as a pseudo-metabolites with the stoichiometric coefficient 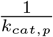, where *k_cat,p_* is the turnover number of protein *p*. The proteins are correspondingly supplied into the network by protein pseudoexchanges. It is in the upper bounds of these pseudoexchanges where the protein concentrations are expressed.

Furthermore, a global resource constraint can be applied to all or a subset of the proteins with unknown concentrations, commonly referred as protein pool.

### Geckopy

A python software package, geckopy, was developed building upon [7]. The package was made open-source under the Apache2.0 license and it can be found at https://github.com/ginkgobioworks/geckopy, documented at https://geckopy.readthedocs.io. The enzyme constrained method stated above is on the core of geckopy, which derives from cobrapy [12] the API and provides an integration layer with pyTFA [10] to apply Thermodynamic Flux Analysis (TFA) on top of the enzyme constrained models.

### Relaxation algorithms

A suite of relaxation algorithms was implemented to compute an Irreducibly Inconsistent Set (IIS); i.e., a minimal set of infeasible constraints. The IIS is resolved by the formulation of different Linear Programming (LP) or Mixed-Integer Linear Programming (MILP) problems that expand the original LP problem ***S*** and may not be unique, therefore a criterion is required to select one set. This corresponds to different objectives of the (MI)LP formulation. As noted in [14], these can be thought as variations of Minimization of Metabolic Adjustment [15]. Three of these relaxation (MI)LPs were implemented:

- Elastic filtering algorithm as detailed in [16]. Briefly, this is a LP problem that introduces elastic variables *e* to a target set of constraints, one for each direction, to recover the feasibility of the solution. For this particular formulation, the target set of constraints are the protein pseudometabolites ***P***, in the upper direction, since the lower bound is always zero. Each constraint vector in **P** is denoted as ***p***. The criterion in this case is the sum minimizing the total flux of the elastic variables e as in 8.

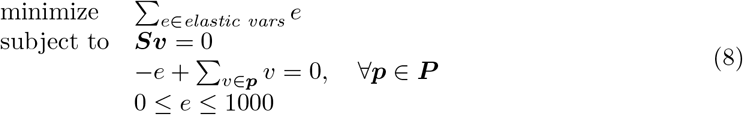 Eq 8 is run iteratively, removing the elastic variables that are found in the previous iteration as part of the optimized objective. This expands the IIS until no elastic variables can be added to obtain a feasible solution. Still, this IIS is not deterministic nor unique and depends on the order at which the iterative sets are retrieved.
- Elastic filtering algorithm as with the previous case, changing the criterion to also optimize the original objective ***Z*** of the problem as in Eq 9.

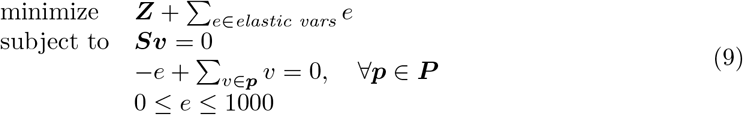
- MILP problem, which minimizes the total count of variables in the IIS. This exploits that the relaxation is only in the upper bounds to avoid quadratic expressions. Thus, two variables are introduced for each protein constraint, an integer variable *i* and linear programming variable *l* as in Eq 10. Multiplying i there is a large constant *K* which is compensated by *l* when needed.

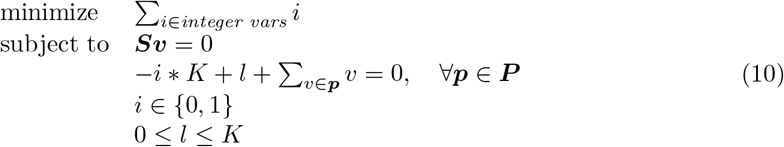

In addition, a greedy brute-force algorithm was provided as it is implemented in Caffeine [17] or in the GECKOmat suite [8]. Briefly, this algorithm removes proteins iteratively guided by the shadow price - i.e., the increment in the objective when one unit of the protein is increased - until the model reaches the growth specified by the user.

The different variants of formulations and objectives reflect different assumptions about the uncertainty of the experimental methods in place. Hence, if it is suspected that the uncertainty is uniformly distributed over all measurements, one of the elastic filtering methods should be chosen over the MILP problem. Correspondingly, if the *a priori* knowledge points to a reduced subset of the proteins with high uncertainty, the MILP might be a better fit. Additionally, the original objective can be included in the relaxation problem. This may be useful when the model is constrained to a known experimental growth rate but the relaxation fails to find a solution with that constraint.

### Proteomics Relaxation benckmark

With the purpose of testing the relaxation in absolute proteomics, the relaxation algorithms implemented in geckopy were benchmarked against a dataset of absolute proteomics across different conditions in *Escherichia coli*, retrieved from [18]. The input of the absolute proteomics datasets, *ϕ*, are defined in Eq 11

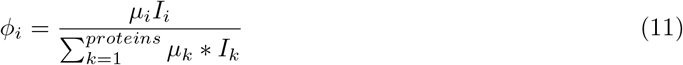

which is the protein mass fraction of each protein *i*,, given the intensity *I_i_* of a quantitative proteomic approach (e.g., xTop, TopPep1/3 and iBAQ) and molecular mass *μ_i_*.

Each *ϕ_p,i_* was converted to the units used by geckopy 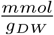, by 12 and 13.

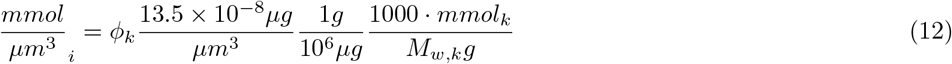

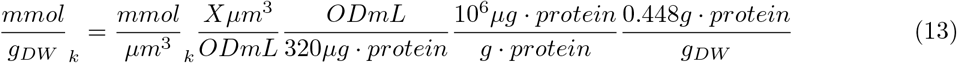

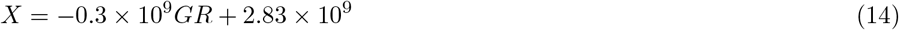

Here, as derived empirically in the supplementary material of [19] 13.5 × 10^-8^ is a constant, *X* is as linear fit of the cellular volume per OD in terms of the growth rate (*GR*, 14) as described ; 320 *μg* is the average number of proteins per *ODmL* for *E. coli* [20]; and *g/g_DW_* was set to 0.448 for all conditions. The latter is an empirical constant although the actual value (or a growth-rate based fit like 14) for the particular experimental conditions would be a more accurate approach.

These constraints were enforced across all 17 conditions, using the upper limit of the Confidence Interval at 95 for concentrations and the lower limit for growth conditions where biological replicates were available. An IIS was solved for all of them individually with the MILP approach, the LP approach, both with and without expanding the IIS. The GEM model of choice was the enzyme constraint version of iML1515 [21], eciML1515 [8]. Just one modification was done to the model, in the case of ammonia limitation, as explained in Table 1.

**Table 1.**
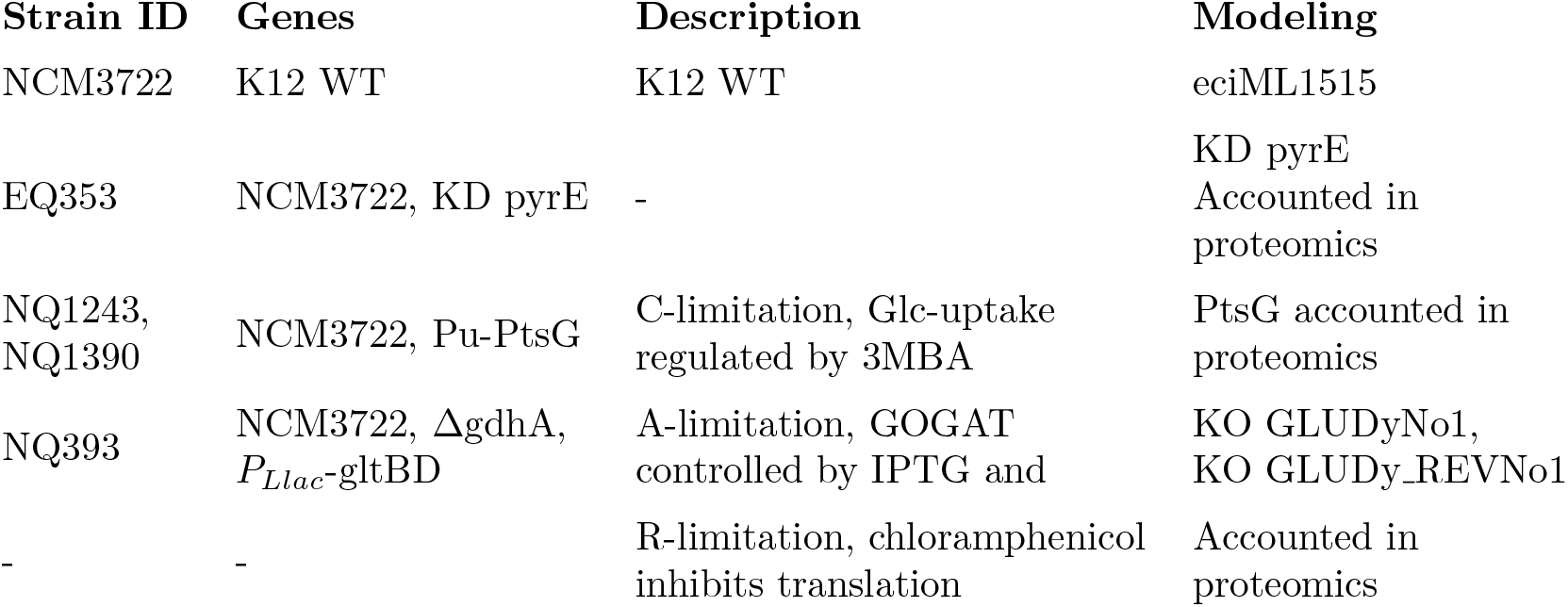
Modeling details of the different strains and conditions for the proteomics datasets of [18]. Here, KD stands for Knock-down. Pu is a *Pseudomonas putida* promoter inserted in Pstd to control it via 3MPA.

The (MI)LP problems were solved using CPLEX [22].

### Flux Variability Analysis comparison

The layered constraints - stoichiometric, thermodymanics, metabolomics, proteomics, and pool assumption - were compared using Flux Variability Analysis (FVA).

FVA is a COBRA algorithm that first runs an FBA problem and fixes the previous objective to its objective value. Then, for each reaction *j*, an FBA problem is formulated where the objective *Z* corresponds to the flux through the *j*; more formally, 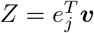. This problem is optimized for both the minimization and maximization directions. The output is the maximum and minimum possible flux of each reaction in the model. Thus, the distribution of maximum and minimum fluxes of the solution space for a particular objective can be inspected using this method.

To ease the interpretation of this analysis, a reduced GEM model was used. Proteins were added to the *Escherichia coli* core model from BiGG [23], with 95 reactions, 72 metabolites and 55 proteins, to form a reduced enzyme constraint model. This model is available at S1 File.

For the sake of a fair comparison, the glucose exchange was fixed to its minimum value when optimizing biomass for all reactions [8]. Although a FVA implementation which block the opposite splitted reaction (common in EC models) to remove spurious variability was implemented as part of geckopy, this model does not have splitted reactions. Moreover, the model does not have isozymes, which further simplifies the analysis. The growth was enforced to be equal to the thermodynamic solution, which did not require relaxation to satisfy a feasible solution.

The relaxation of experimental data used was always the IIS MILP (Eq 10) + the greedy filter over the IIS, which minimizes the number of constraints removed from the problem.

### Average enzyme saturation

The average enzyme saturation was calculated as implemented in geckopy, following Eq 15

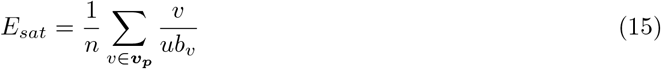

where ***v_p_*** is the vector of protein exchange variables whose upper bound in non zero, *ub_v_* is the upper bound of the given variable; and *n* is the length of ***v_p_***.

## Results

### Geckopy

#### Design principles

Three design principles were chosen for the development of geckopy:

1. SBML compliance, to maintain the community standards.
2. API separation of the proteins from metabolites and reactions.
3. Ease of experimental data loading.

#### SBML specification

As in previous iterations of enzyme constraint models [7,8], proteins are *SBML Species* [24]. This change moves away from naming conventions to identify proteins with a structured system that more appropriately complies with the FAIR principles [25]. Here, we proposed to extend this definition to proteins as *Species*:

- in a defined SBML Group [4] named ”Protein”;
- with the concentration specified as the optional Species attribute *initialAmount*; and
- whose *k_cat_* value is specified in the stoichiometry coefficient of the reactions, as usual.

The reasoning behind this specification is to ensure that the document is self-contained and does not depend on naming conventions during the parsing. Thus, modelers can save in the document proteomics information if needed and the SBML standard is preserved.

**Listing 1.**
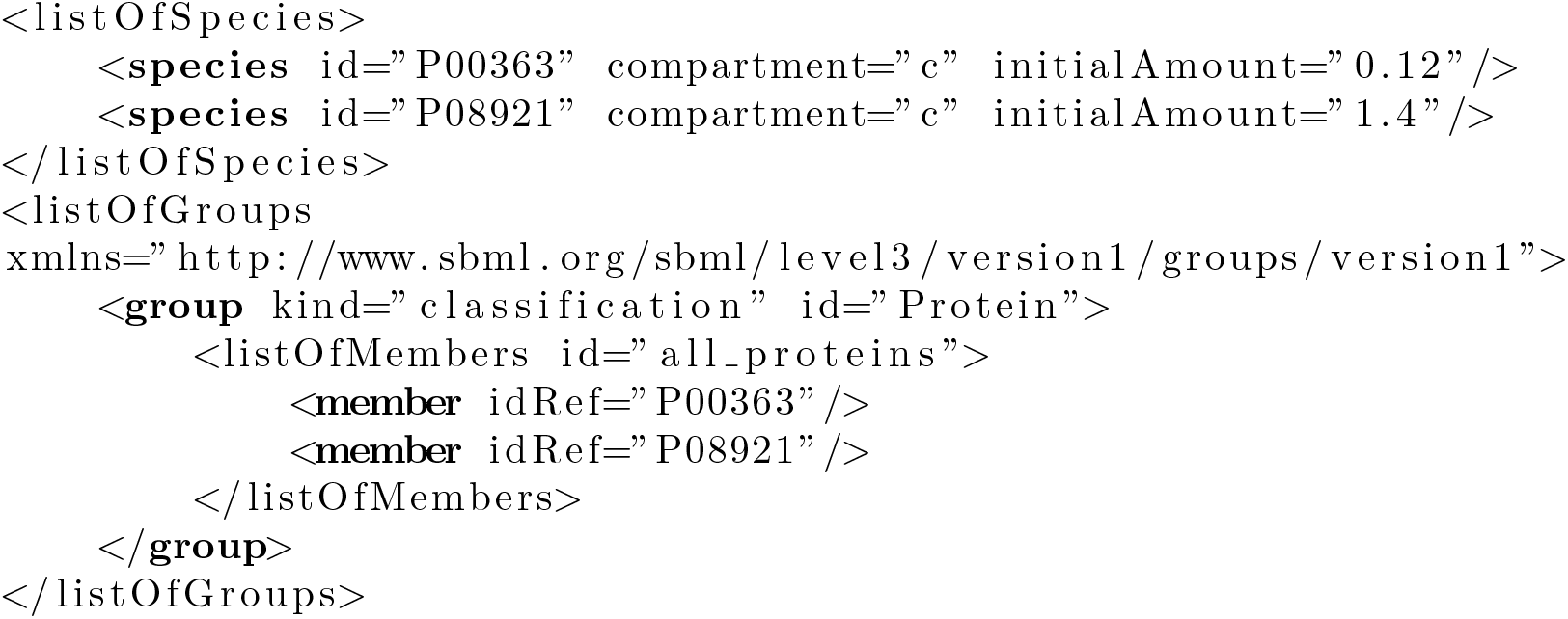
SBML document with proteins.

Geckopy implements the GECKO algorithm [7]. Enzyme constraint models expand the stoichiometry matrix of the classic Flux Balance Analysis formulation [2] - designated as ***S*** - with proteins constraints - designated as ***P***. The proteins are supplied by variables with an upper bound that reflect the experimental measure when known, i.e., pseudoexchanges that supply the protein to be consumed in the reactions; and appear as constraints in the reaction variables where they participate, i.e., pseudometabolites with a stoichiometric coefficient as the inverse *k_cat_*. Geckopy implements this, built on top the established COBRA software cobrapy [12]. Additionally, geckopy includes a suite of flux analysis algorithms for enzyme constraint models analogous to the ones in cobrapy, such as Flux Variability Analysis, parsimonious Flux Protein Analysis, etc.

Another difference with the GECKO toolbox in MATLAB [8] is that geckopy implements the reversible reactions internally, removing the need of splitting all of the reversible reactions in the SBML. Nonetheless, geckopy is able to read *legacy* enzyme constraint models that use splitted reactions and/or proteins specified with the naming convention *prot_UNIPROTID.*

### Benchmark of relaxation algorithms with proteomics datasets

The relaxation of experimental constraints is usually a requirement to yield a feasible enzyme constraint model. In order to test the assumptions and show the utility of the relaxation algorithms suite, they were benchmarked against a dataset of *Escherichia coli* growing in 17 conditions (carbon, ammonia and ribosome limitation) from [18].

For each condition, the eciML1515 model was constrained with the reported protein measurements and the experimental growth rate (see Methods). Of the 1259 proteins in the model, 1243 were found in the experimental data consistently across conditions. The number of conditions where a protein concentration had to be relaxed is shown in Fig 1. Across samples, the average number of relaxed proteins was 94.94 and 95.17 for the MILP and MILP first IIS’s, respectively, with standard deviations of 10.57 and 11.21. The number of relaxed proteins in the LP and LP first cases, with means of 74.23 and 75.82 and standard deviations of 11.54 and 12.82, respectively. In contrast, the average enzyme saturation in Fig 2 for the LP first and the MILP first methods were similar across samples (below the 20%), but with an overall decrease for the MILP case.

For the LP cases, inspecting the relaxation values of the proteins - i.e., how much the experimental measurement must be violated - is interesting to assess possible errors in the model or the experimental data. This informs not only about a protein being relaxed, but about how much it was relaxed. A closer look at the relaxation values in the LP first case in Fig 4 revealed that the number of proteins in the IIS drops down to 74.05 when taking into account only primal values greater than 10^-7^ (which is on the threshold for numerical errors). Moreover, 10 proteins hold around 77.12% and 90.97% of the total relaxation of the problem across the samples. These proteins were the Phosphoribosylformylglycinamidine synthase, the Aconitase, subunits of the ATP synthase and the 3-oxoacyl-[acyl-carrier-protein] synthase 1. One deviation form this trend was the R-limitation positive control, which additionally showed two proteins with a large relaxation value: a Phosphoserine phosphatase and a Riboflavin reductase.

As expected, it should be noted in Fig 4 that the growth is highly correlated - Pearson’s correlation coefficient of 0.966 (Fig 3) - with the sum of the relaxation values. This indicates that the protein constraints are less important when moved away from optimal conditions, whereas uptake rates dominate the solution in these cases. Once in optimal conditions, the higher relaxation values may be highlighting an overall underestimation of *k_cat_* values that was not observed in lower growth rates, when operating under non-saturation rates.

### Layered comparison of constraints

Geckopy implements an integration API with pytfa [10] to add thermodynamic and metabolomics constraints on top of the enzyme-constraint models. Thus, it was relevant to check if the solution space becomes smaller - and thus less uncertain - when applying combinations of these different constraints.

The comparison of FVA runs shows the effect of the different constraints layered in the reduction of the solution space in Fig 5. The relaxation for the ”Thermodynamics + Proteomics” scenario removed 22 proteins; and the ”Proteomics” removed 18. It can be observed in Fig 5A that the flux interval distribution for the proteomics constraint is the less dispersed, followed by the thermo and proteomics constraints, the thermodynamic constraints, the pool constraint and, finally, the flux balance analysis. This is reflected in Fig 5B, where it can be seen how the different constraints can turn a reversible reaction into irreversible (TPI, proteomics), reduce the flux allowed, fix it to a reduced range (PPS, proteomics) or block it completely (PPCK).

## Discussion

### The relaxation of the enzyme constrain model reveals inconsistencies between the model and the experimental measurements

The exploration of the IIS of several proteomics datasets revealed 10 proteins that are significantly and consistently clashing between the model and the data. Of these proteins, 6 are subunits of the ATPase, which is not only a transmembrane protein - which can pose quantification challenges to proteomics methods - but also a protein complex with complex kinetics where the enzyme constraint assumption may not hold as well as with the usual metabolic enzymes, since a *k_cat_* cannot summarise the mechanochemical operation of the ATPase [26]. Among the other four, we found *k_cat_* values in the first percentile of the *k_cat_* values in the model, indicating possible underestimations of the actual *in vivo k_cat_* values by *in vitro* methods.

This information, consistent across samples and experimental methods, could be used to flag regions where the protein coverage of the model can be improved, kinetic parameters updated, or where experimental methods may lead to measurements errors. There were two outliers in the R-limitation (translation inhibition) positive control condition: the phosphoserine phosphatase and one isozyme of the FMN reductase. These two enzymes also have comparably low *k_cat_* values and are the responsible for the total relaxation being higher than expected for the growth rate of the condition (most right point in 3). It is important to note that this condition exhibited the highest growth rate, exceeding the growth of the model, so it is possible that the experimental Glc uptake rate also exceeded the modeled value of 10 *mmol/gDW/h*, impacting the relaxation.

Overall, the number of relaxed enzyme contraints in the MILP problems was greater than the LP counterparts overall, producing less constrained problems as represented by the decrease in average enzyme saturation. Therefore, it may be prefereable to choose the LP elastic relaxation instead, which only adjust the experimental measurements, instead of the MILP, which removes them. Moreover, the LP formulation has the advantage of providing relaxation values for each relaxed protein, providing a measure of the extent of the violation of the experimental data.

### Layered comparisons of constraints reveals inconsistencies when combining experimental constraints of different nature

As it was reported in 5, enzyme constraints result in a more constrained space solution than the thermodynamic constraints. This was expected, as 47 reactions (49.47%) in the core model are already unidirectional, which is the main source of constraints that TFA is expected to enforce [9].

It was less expected that the proteomics constraints alone would produce a smaller solution space than the enzyme constraints + thermodynamics. However, this is clearly explained by looking at the number of proteins removed from the problem in the two cases. This indicates that absolute proteomics constraints introduce inconsistencies between the experimental measurements and the model which may arise from a) inaccurate experimental measurements b) lack of coverage of isozymes and promiscuity in the model c) lack of coverage of reactions in the model. In this case, the model presented is a reduced reconstruction, which is likely the greater source of inconsistencies.

It is worth noting that this behavior of an increased solution space for combinations of thermodynamics and enzyme constraints had been previously reported [14]. Furthermore, better management of uncertainty and the use of more accurate methods to estimate free energies, like component contribution, could provide more informative thermodynamics constraints that enhance the predictions, as in [27].

## Conclusion

In this work, we have presented Geckopy 3.0 a package that, beyond the enzyme-constraint formulation provides three innovations:

1. A way to include Proteins as first class citizens in the SBML document, complying with the community standards, with its corresponding parser implementation.
2. Inspection and biological discussion of a suite of relaxation algorithms, inspired by [14], which may be used to reveal inconsistencies in the model or in the proteomics data if used systematically.
3. An integration with pytfa [10] to include thermodynamic and metabolomics constraints on top of the enzyme constraints. The inspection of this integration revealed that combining both proteomics and metabolomics may not provide a more constraint space than one of them alone, in accordance with previous works in the field [14].

In future work, the inclusion of multiTFA [27] instead of pytfa and more sophisticated approaches to combine both enzyme-constraint and proteomics constraints might be explored. Overall, we would like to emphasize the need in the field to integrate multi-omics methods in a consistent manner.

## Supporting information

**S1 File. Reduced *E. coli* genome-scale network reconstruction.** The model contains 95 reactions, 72 metabolites and 55 proteins.

**S2 File. Relaxation noteboook.** Jupyter notebook to reproduce Figures 1, 2, 3, 4.

**S3 File. FVA noteboook.** Jupyter notebook to reproduce Figure 5.

## Acknowledgments

We would like to acknowledge Benjamín Sánchez and Iván Domenzáin for their valuable perspectives and discussions on enzyme-constraint models. We would also like to thank the Ginkgo Bioworks team who kindly spent time hearing about geckopy during its development and offering feedback. N.S. acknowledges funding from the Novo Nordisk Foundation under the Fermentation Based Biomanufacturing program (grant no. NNF17SA0031362).

