## Supplementary material for "Simultaneous application of enzyme and thermodynamic constraints to metabolic models using an updated Python implementation of GECKO": S2 FIle: proteomics_data_relaxations.ipynb

```
# %% [markdown]
# Proteomics EC GEM integration

#### Table of contents
* [Data preparation](#Data-preparation)
    * [Uniprot Mapping](#Loading-the-data)
    * [Protein unit conversion](#Protein-unit-conversion)
* [Application to model](#Application-to-model)
* [Relaxation runs](#Relaxation-runs)
    * [Relaxation Aggregated by biological replicates](#Relaxation-Aggregated-by-biological-replicates)
    * [Sensibility Aggregated by biological replicates](#Sensibility-Aggregated-by-biological-replicates)
    * [Get average enzyme saturation](#Get-average-enzyme-saturation)

# %% [code]
import re
import warnings
from collections import Counter
from copy import deepcopy
from functools import reduce
from math import isnan, sqrt
from pathlib import Path

import geckopy
import numpy as np
import pandas as pd
import plotly.express as px
from geckopy.experimental import from_mmol_gDW
from geckopy.experimental.molecular_weights import extract_proteins
from geckopy.experimental.relaxation import (
    Objective_rule,
    elastic_upper_relaxation,
    get_upper_relaxation,
)
from tqdm import tqdm

# %% [markdown]
## Data preparation

# %% [markdown]
### Uniprot extraction

# %% [code]
DATA = Path.cwd().parent / "data"

# %% [code]
ev9 = pd.read_excel(
    DATA / "msb20209536-sup-0010-datasetev9.xlsx",
    engine="openpyxl",
    sheet_name=1,
)

# %% [code]
print(
    pd.read_excel(
        DATA / "msb20209536-sup-0010-datasetev9.xlsx", engine="openpyxl", sheet_name=0
    ).iloc[1, 0]
)

# %% [code]
ev9.describe()

# %% [code]
PROT_PATTERN = re.compile(
    r"[OPQ][0-9][A-Z0-9]{3}[0-9]|[A-NR-Z][0-9]([A-Z][A-Z0-9]{2}[0-9]){1,2}"
)

# %% [code]
ev9["Protein ID"].isna().sum() / ev9.shape[0]

# %% [markdown]
2.7% of proteins do not have an identifier. We can just remove them for now.

# %% [code]
ev9_valid = ev9[~ev9["Protein ID"].isna()]

# %% [code]
ev9_valid["Protein ID"].str.match(PROT_PATTERN).all()

# %% [markdown]
All proteins seems to match a Uniprot ID.

# %% [markdown]
In preparation for the unit conversions, extract the molecular weights querying uniprot.

# %% [code]
prot_list = ev9_valid["Protein ID"].to_list()
prot_reactions = [f"prot_{prot}" for prot in prot_list]
prot_dict = dict(zip(prot_list, prot_reactions))

# %% [code]
ev9_valid["MW"] = extract_proteins(None, prot_dict)["MW"]

# %% [code]
ev9_valid = ev9_valid[~ev9_valid["MW"].isna()]
ev9_valid.to_csv(DATA / "EV9_clean_MW.tsv", sep="\t")

# %% [markdown]
### Protein unit conversion

# %% [code]
ev9_valid = pd.read_csv(DATA / "EV9_clean_MW.tsv", sep="\t")

# %% [code]
ev9_valid = ev9_valid.loc[:, ~ev9_valid.columns.str.startswith("Unnamed")]

# %% [markdown]
Let's put it in tidy format to make the unit conversions.

# %% [code]
ev9_tidy = ev9_valid.melt(
    ["MW", "Protein ID", "Gene name", "Gene locus"],
    var_name="Sample",
    value_name="Mass_fraction",
)

# %% [code]
g_per_gDW = 0.448

# %% [markdown]
\begin{align}
\frac{\mu g}{\mu m^3}_k &= \phi_k 13.5 x 10^{-8} \\
v_{ODmL} &= \frac{X \mu m^3}{OD mL} \\ 
\frac{mmol}{g_{DW}}_k &= \frac{k \mu g}{\mu m^3}_k \frac{1 g}{10^6 \mu g} \frac{1000 mmol_k}{M_{w,k} g} v_{ODmL} \frac{OD mL}{320 \mu g protein}  \frac{10^6 \mu g \text{protein}}{g \text{protein}} \frac{0.448g protein}{g_{DW}}
\end{align}


# %% [code]
growth_conditions = pd.read_excel(
    DATA / "msb20209536-sup-0004-datasetev3.xlsx", sheet_name=1, engine="openpyxl"
)

# %% [code]
sample_growth = growth_conditions[["Sample ID", "Growth rate (1/h)"]].set_index(
    "Sample ID"
)["Growth rate (1/h)"]

# %% [code]
ev9_v_ODmL = ev9_tidy.Sample.apply(lambda x: sample_growth.loc[x] * -0.3e9 + 2.83e9)

# %% [code]
ev9_tidy["mmolgDW"] = (
    13.5e-8 * ev9_tidy.Mass_fraction * 1000 / ev9_tidy.MW * ev9_v_ODmL / 320 * g_per_gDW
)

# %% [code]
ev9_tidy.mean()

# %% [code]
ev9_tidy[["Protein ID", "mmolgDW", "MW", "Sample"]].to_csv(
    DATA / "EV9_final.tsv", sep="\t", index=False
)

# %% [code]
ev9_final = pd.read_csv(DATA / "EV9_final.tsv", sep="\t")

# %% [code]
ev9_final = ev9_final[~ev9_final.mmolgDW.isna()]

# %% [code]
df_prot = ev9_final

# %% [code]
df_prot.to_csv(DATA / "processed_proteins.tsv", sep="\t", index=False)

# %% [markdown]
### Application to model

# %% [markdown]
The absolute proteomics will be applied to the model for every condition.

# %% [code]
df_prot = pd.read_csv(DATA / "processed_proteins.tsv", sep="\t")

# %% [code]
df_prot.iloc[[0, 1, 2, 3, -5, -4, -3, -2, -1], :]

# %% [code]
model = geckopy.io.read_sbml_ec_model(DATA / "eciML1515.xml.gz")

# %% [code]
model.solver = "cplex"

# %% [code]
growth_conditions = pd.read_excel(
    DATA / "msb20209536-sup-0004-datasetev3.xlsx", sheet_name=1, engine="openpyxl"
)

# %% [code]
growth_conditions.head()

# %% [markdown]
Need the growth rate $\frac{1}{h}$ (in one of the sheets).

# %% [code]
growth_conditions.columns

# %% [code]
growth_conditions.loc[growth_conditions["Sample ID"] == "A1-1", "Growth rate (1/h)"][0]

# %% [code]
objective_id = "BIOMASS_Ec_iML1515_core_75p37M"

# %% [code]
iis_lp_counter = dict()
iis_milp_counter = dict()

# %% [code]
growth_conditions["Sample ID"]

# %% [code]
growth_conditions.loc[
    growth_conditions["Sample ID"] == "A1-3", "Growth rate (1/h)"
].iloc[0]

# %% [markdown]
# Relaxation runs

# %% [markdown]
## Relaxation Aggregated by biological replicates
Lastly, do it properly with the biological replicates (but different original datasets).

# %% [markdown]
#### About strains
EQ353 is equivalent to the model:

- Knock down in pyrE expression (corresponds to ORPTNo1, prot_P0A7E3, which should be accounted in the proteomics data).
- Knock out of ilvG, which is already removed in the model.

NCM3722 is the model (K12 usual strain).
- <span style="color:#e26b73">Is $\Delta$ilvG also in the original K12?</span>

NQ1243, NQ1390 (C-limitation) both have the titration construct Pu-ptsG that regulates Glc uptake externally using 3MBA (no difference is specified between those, no strains with those names are in the specified references).

- Can we expect this from proteomics?

NQ393 (A-limitation)
- <span style="color:#e26b73">Deletion of $\Delta$gdhA, GDH: H2O + L-glutamate + NADP+ = 2-oxoglutarate + H+ + NADPH + NH4+*</span>.
- Promoter replacement of gltBD to control (via IPTG) GOGAT expression: Gln + akg -> 2 Glu (should be accounted for in proteomics).

#### About the growth conditions
C-limitation (carbon)
- 3MBA supplement, which is proportional to ptsG expression (Glc uptake).

A-limitation (ammonia)
- IPTG supplement, which is proportional to GOGAT expression.

R-limitation (ribosome)
- Cloramphenicol supplements, which inhibit ribosomal translation (it is seen in a cap of the growth rate).

# %% [code]
# so that groupby makes sense
growth_conditions.at[growth_conditions.Supplement.isna(), "Supplement"] = "No"

# %% [code]
(
    growth_conditions.groupby(["Strain", "Supplement"]).agg(
        {
            "Growth rate (1/h)": lambda x: x.mean() if len(x) > 1 else x.iloc[0],
            "Sample ID": lambda x: x.to_list(),
        }
    )
)

# %% [code]
df_growth_agg_changes = growth_conditions.groupby(["Strain", "Supplement"]).agg(
    growth_mean=pd.NamedAgg(column="Growth rate (1/h)", aggfunc="mean"),
    growth_std=pd.NamedAgg(column="Growth rate (1/h)", aggfunc="std"),
    sample=pd.NamedAgg(column="Sample ID", aggfunc=lambda x: x.to_list()),
)

# %% [code]
iis_lp_dict_agg = {}
iis_milp_dict_agg = {}
iis_lp_dict_agg_first = {}
iis_milp_dict_agg_first = {}

# %% [code]
df_growth_agg_changes

# %% [markdown]
The protein identifiers in the model start with "prot_".

# %% [code]
df_prot["model_id"] = df_prot["Protein ID"].apply(lambda prot: f"prot_{prot}")

# %% [markdown]
Mark the proteins in the model to filter them during the condition selection.

This is not required (geckopy will ignore the unknown proteins) but will remove a lot of the warnings when applying protein concentrations of `from_mmol_gDW`.

# %% [code]
df_prot["in_model"] = df_prot["model_id"].apply(lambda prot: prot in model.proteins)

# %% [code]
df_prot.to_csv(DATA / "processed_proteins.tsv", sep="\t", index=False)

# %% [markdown]
In addition to the number of proteins per sample per method, the primal (relaxation) values will be gathered in a dataframe for the LP first case to check how those are distributed and how numerical errors affect the solution.

# %% [code]
df_results = pd.DataFrame({"sample": [], "protein": [], "primal": [], "dual": []})
df_results

# %% [markdown]
For the conditions that exceed model unconstrained growth, the report growth will be lowered to the unconstrained maximum growth. Likely, the glucose uptake is greater than the one in the model but we do not have information about it.

# %% [code]
unconstrained_growth = model.slim_optimize()

# %% [code]
for i, (growth_mean, growth_std, sample) in tqdm(
    df_growth_agg_changes[["growth_mean", "growth_std", "sample"]].iterrows(),
    total=df_growth_agg_changes[["growth_mean", "growth_std", "sample"]].shape[0],
    desc="Agg samples IIS",
):
    if len(sample) > 1:
        df_sample = (
            df_prot.loc[df_prot.Sample.isin(sample) & df_prot.in_model, ["model_id", "mmolgDW"]]
            .groupby("model_id")
            .agg([np.mean, np.std, "count"])
            .droplevel(0, axis=1)
            # use the upper limit of the CI at 0.95
            .assign(
                mmolgDW=lambda x: x["mean"] + 1.960 * x["std"] / np.sqrt(x["count"] - 1)
            )
            .reset_index()
        )
        # use the lower limit of the CI at 0.95
        growth = growth_mean - 1.960 * growth_std / sqrt(len(sample) - 1)
    else:
        df_sample = df_prot.loc[df_prot.Sample.isin(sample)  & df_prot.in_model, ["model_id", "mmolgDW"]]
        growth = growth_mean
    df_sample.index = df_sample["model_id"]
    # apply concentrations (this is not inplace)
    with warnings.catch_warnings():
        warnings.simplefilter("ignore")
        model_conc = from_mmol_gDW(model, df_sample["mmolgDW"])
    # force a growth condition before relaxing
    model_conc.reactions.get_by_id(objective_id).lower_bound = (
        growth if growth < unconstrained_growth else unconstrained_growth
    )
    prot_candidates = [
        prot.id for prot in model_conc.proteins if prot.concentration is not None
    ]
    prot_candidates = [
        prot_id
        for prot_id in prot_candidates
        if not isnan(model_conc.proteins.get_by_id(prot_id).concentration)
    ]
    iis_lp_agg = elastic_upper_relaxation(
        model_conc, prot_candidates, Objective_rule.MIN_ELASTIC_SUM
    )
    iis_milp_agg = elastic_upper_relaxation(
        model_conc, prot_candidates, Objective_rule.MIN_MILP_COUNT
    )
    iis_milp_agg_first, milp_status = get_upper_relaxation(
        model_conc.copy(), prot_candidates, Objective_rule.MIN_MILP_COUNT
    )
    iis_lp_agg_first, lp_status = get_upper_relaxation(
        model_conc, prot_candidates, Objective_rule.MIN_ELASTIC_SUM
    )
    iis_lp_dict_agg[str(sample)] = iis_lp_agg
    iis_milp_dict_agg[str(sample)] = iis_milp_agg
    iis_lp_dict_agg_first[str(sample)] = iis_lp_agg_first
    iis_milp_dict_agg_first[str(sample)] = iis_milp_agg_first
    # extract info for this sample
    if lp_status == "optimal":
        df_results = pd.concat(
            [
                df_results,
                pd.DataFrame(
                    {
                        "sample": len(iis_lp_agg_first) * [str(sample)],
                        "protein": [prot_id for prot_id in iis_lp_agg_first],
                        "primal": [
                            model_conc.variables.get(f"V_{prot_id}").primal
                            for prot_id in iis_lp_agg_first
                        ],
                        "dual": [
                            model_conc.variables.get(f"V_{prot_id}").dual
                            for prot_id in iis_lp_agg_first
                        ],
                        "method": "LP first"
                    }
                ),
            ]
        )

# %% [code]
tidy_results_agg = pd.DataFrame(
    {
        "protein": reduce(lambda a, b: list(a) + list(b), iis_lp_dict_agg.values(), []),
        "sample": reduce(
            lambda a, b: a + [b[0]] * len(b[1]), iis_lp_dict_agg.items(), []
        ),
    }
)
tidy_results_agg["method"] = "LP"
milp_results_agg = pd.DataFrame(
    {
        "protein": reduce(
            lambda a, b: list(a) + list(b), iis_milp_dict_agg.values(), []
        ),
        "sample": reduce(
            lambda a, b: a + [b[0]] * len(b[1]), iis_milp_dict_agg.items(), []
        ),
    }
)
milp_results_agg["method"] = "MILP"
tidy_results_agg_first = pd.DataFrame(
    {
        "protein": reduce(
            lambda a, b: list(a) + list(b), iis_lp_dict_agg_first.values(), []
        ),
        "sample": reduce(
            lambda a, b: a + [b[0]] * len(b[1]), iis_lp_dict_agg_first.items(), []
        ),
    }
)
tidy_results_agg_first["method"] = "LP first"
milp_results_agg_first = pd.DataFrame(
    {
        "protein": reduce(
            lambda a, b: list(a) + list(b), iis_milp_dict_agg_first.values(), []
        ),
        "sample": reduce(
            lambda a, b: a + [b[0]] * len(b[1]), iis_milp_dict_agg_first.items(), []
        ),
    }
)
milp_results_agg_first["method"] = "MILP first"
final_results_agg = pd.concat(
    [
        tidy_results_agg,
        milp_results_agg,
        tidy_results_agg_first,
        milp_results_agg_first,
    ],
    axis=0,
)

# %% [code]
final_results_agg

# %% [code]
final_results_agg.to_csv(
    DATA / "results_merged_allLP_AGG_wfirst.tsv", sep="\t", index=False
)

# %% [code]
final_results_agg = pd.read_csv(DATA / "results_merged_allLP_AGG_wfirst.tsv", sep="\t")

# %% [code]
fig = px.histogram(
    final_results_agg.groupby(["protein", "method"])
    .count()
    .reset_index(drop=False)
    .rename({"sample": "appearances in samples"}, axis=1),
    x="appearances in samples",
    color="method",
    barmode="group",
    marginal="box",
    width=800,
    height=400,
    template="plotly_white",
)
fig.update_layout(
    legend=dict(
        x=0.1,
        y=0.5,
        traceorder="normal",
        font=dict(family="sans-serif", size=15, color="black"),
    )
)
fig.update_xaxes(titlefont=dict(size=21))
fig.update_yaxes(titlefont=dict(size=21))
fig

# %% [code]
515 / 1259

# %% [code]
final_results_agg.groupby(["sample", "method"]).count().reset_index(drop=False).groupby(
    "method"
).agg([np.mean, np.std])

# %% [code]
fig.write_image("results_0448_relax_proteo_mori2021_rep1.pdf")

# %% [code]
set_prots = list(iis_milp_dict_agg_first.values())[0:2]

# %% [code]
len(set_prots)

# %% [markdown]
#### Sensibility Aggregated by biological replicates

Generate the plot to inspect the relaxation on the LP first problem.

# %% [code]
# save results
df_results.to_csv(
    DATA / "results_merged_LP_first_AGG_sens_with_cap.tsv", sep="\t", index=False
)

# %% [code]
df_results = pd.read_csv(
    DATA / "results_merged_LP_first_AGG_sens_with_cap.tsv", sep="\t"
)

# %% [code]
def max_n(group):
    return group.loc[
        group["primal"] >= group["primal"].sort_values(ascending=False).iloc[10]
    ].sort_values("primal", ascending=False)

def more_than(group, than=1e-7):
    return group.loc[group["primal"] >= than].sort_values("primal", ascending=False)

# %% [code]
df_results_top10_sample = (
    df_results.groupby("sample").apply(max_n).reset_index(drop=True)
)

# %% [code]
fig_lp_sample_more_than = px.bar(
    df_results_top10_sample,
    color="protein",
    y="primal",
    x="sample",
    template="plotly_white",
)

fig_lp_sample_more_than

# %% [code]
df_results_top10_sample = (
    df_results.groupby("sample").apply(max_n).reset_index(drop=True)
)
df_results_top10_sample["sample"] = df_results_top10_sample["sample"].replace(
    {s: i for i, s in enumerate(df_results_top10_sample["sample"].unique())}
)

# %% [code]
df_results_top10_sample

# %% [code]
fig_lp_sample_more_than = px.bar(
    df_results_top10_sample,
    color="protein",
    y="primal",
    x="sample",
    template="plotly_white",
)

fig_lp_sample_more_than

# %% [code]
fig_lp_sample_more_than.write_image(
    "results_0448_relax_proteo_mori2021_sample_more_than_lpfirst.pdf"
)

# %% [code]
model.reactions.get_by_id("PRFGSNo1")

# %% [code]
model.proteins.prot_P0ABA6.kcats

# %% [code]
{
    prot: [reac.name for reac in model.proteins.get_by_id(prot).reactions]
    for prot in df_results_top10_sample.loc[
        df_results_top10_sample["sample"] == 20, "protein"
    ].to_list()
}

# %% [code]
df_results_more_than_sample = (
    df_results.groupby("sample")
    .apply(lambda x: more_than(x, 1e-7))
    .reset_index(drop=True)
)

# %% [code]
df_results_more_than_sample.groupby("sample").count().mean()

# %% [code]
# df_results_more_than_sample["Sample (number)"] = df_results_more_than_sample["sample"].replace(
#     {s: i for i, s in enumerate(df_results_more_than_sample["sample"].unique())}
# )

# %% [code]
df_growth_agg_changes

# %% [code]
ticktext = [
    "Reference EQ353",
    "C-limitation (400 µM 3MBA) NQ1243",
    "R-limitation NCM3722",
    "C-limitation (100 µM 3MBA) NQ1243",
    "C-limitation NQ1243",
    "C-limitation (400 µM 3MBA) NQ1390",
    "C-limitation (40 µM 3MBA) NQ1390",
    "A-limitation (20 µM IPTG) NQ393",
    "A-limitation (30 µM IPTG) NQ393",
    "A-limitation (40 µM IPTG) NQ393",
    "A-limitation (50 µM IPTG) NQ393",
    "A-limitation (100 µM IPTG) NQ393",
    "C-limitation (400 µg 3MBA) NQ1390",
    "R-limitation (2µg CAF) NCM3722",
    "R-limitation (4µg CAF) NCM3722",
    "R-limitation (6µg CAF) NCM3722",
    "R-limitation (8µg CAF) NCM3722",
]

# %% [code]
tickvals = [
    "['A1-1', 'A1-2', 'A1-3', 'C1', 'F1-1', 'F1-2', 'F1-3']",
    "['C3', 'F4']",
    "['C2', 'A2', 'H1', 'H5']",
    "['C4', 'C8', 'F5']",
    "['C5', 'D6', 'F6']",
    "['C6', 'F7']",
    "['C7', 'D7', 'F8']",
    "['D1']",
    "['D2']",
    "['D3']",
    "['D4']",
    "['D5', 'F2', 'F3']",
    "['D8']",
    "['E1']",
    "['E2']",
    "['E3']",
    "['E4']",
]

# %% [code]
print(
    f"Relaxation results: {len(df_results_more_than_sample['sample'].unique())}\n"
    f"Ticktext len: {len(ticktext)}\n"
    f"Tickvals len: {len(tickvals)}\n"
)

# %% [code]
fig = px.bar(
    df_results_more_than_sample,
    color="protein",
    y="primal",
    x="sample",
    hover_data=["sample"],
    width=1200,
    height=800,
    template="plotly_white",
)

fig.update_layout(
    xaxis={"title": "Conditions"},
    yaxis={"title": "relaxation primal value (> 1e-7)"},
)

fig.update_xaxes(
    ticktext=ticktext,
    tickvals=tickvals,
)

fig.update_xaxes(titlefont=dict(size=21), tickfont=dict(size=16), tickangle=50)
fig.update_yaxes(titlefont=dict(size=21))

# %% [code]
model.proteins.prot_P0AGB0.kcats["PSP_LNo1"]

# %% [code]
model.reactions.PSP_LNo1

# %% [code]
sum(model.proteins.prot_P0AEN1.kcats.values()) / len(
    model.proteins.prot_P0AEN1.kcats.items()
)

# %% [code]
model.reactions.FMNRxNo2

# %% [code]
fig.write_image("results_0448_relax_proteo_mori2021_sample_more_than_lpfirst.pdf")

# %% [markdown]
Linear fit of total relaxation ~ growth rate.

# %% [code]
growth_total_primal = pd.merge(
    df_results_more_than_sample.groupby("sample").sum().reset_index(),
    df_growth_agg_changes[["sample", "growth_mean"]].assign(
        sample=df_growth_agg_changes["sample"].astype(str)
    ),
    on="sample",
)

# %% [code]
x = growth_total_primal.primal.to_numpy()
y = growth_total_primal.growth_mean.to_numpy()

# %% [code]
y

# %% [code]
from scipy.stats import pearsonr

# %% [code]
pearsonr(x, y)

# %% [code]
m, c = np.linalg.lstsq(np.vstack([x, np.ones(len(x))]).T, y, rcond=None)[0]

# %% [code]
import matplotlib.pyplot as plt

plt.rcParams["figure.figsize"] = (16, 10)
plt.rcParams.update({"font.size": 22})

# %% [code]
plt.style.use("seaborn-whitegrid")

# %% [code]
unconstrained_growth = model.slim_optimize()

# %% [code]
_ = plt.plot(
    x[y > unconstrained_growth],
    y[y > unconstrained_growth],
    "s",
    color="#00cc96",
    label="Exceeding model growth",
    markersize=15,
)
_ = plt.plot(x, y, "o", color="#636efa", label="Original data", markersize=10)

_ = plt.plot(x, m * x + c, color="#ef553b", label="Fitted line")
plt.xlabel("Sum of relaxation")
plt.ylabel("Growth rate")
_ = plt.legend(frameon=True, edgecolor="white")

plt.savefig("fitting_growth_relaxation.png", dpi=300)
plt.show()

# %% [code]
df_results_more_than_sample.groupby("sample").count()["protein"].agg(["mean", "std"])

# %% [code]
df_results_more_than6_sample = (
    df_results.groupby("sample")
    .apply(lambda x: more_than(x, 1e-6))
    .reset_index(drop=True)
)

# %% [code]
fig = px.histogram(
    df_results_more_than6_sample.groupby(["protein"]).count().reset_index(drop=False),
    x="sample",
    barmode="group",
    marginal="box",
    width=800,
    height=400,
    template="plotly_white",
)
fig.update_layout(
    legend=dict(
        x=0.1,
        y=0.5,
        traceorder="normal",
        font=dict(family="sans-serif", size=12, color="black"),
    )
)
fig

# %% [markdown]
## Get average enzyme saturation

# %% [code]
from geckopy.flux_analysis import (
    count_protein_saturation,
    get_average_protein_saturation,
)

# %% [code]
growth_conditions = pd.read_excel(
    DATA / "msb20209536-sup-0004-datasetev3.xlsx", sheet_name=1, engine="openpyxl"
)
growth_conditions.at[growth_conditions.Supplement.isna(), "Supplement"] = "No"
# group growth conditions by biological replicates
df_growth_agg_changes = growth_conditions.groupby(["Strain", "Supplement"]).agg(
    growth_mean=pd.NamedAgg(column="Growth rate (1/h)", aggfunc="mean"),
    growth_std=pd.NamedAgg(column="Growth rate (1/h)", aggfunc="std"),
    sample=pd.NamedAgg(column="Sample ID", aggfunc=lambda x: x.to_list()),
)
# add info to later delete gldha on NQ393
df_growth_agg_changes["gdha_del"] = (
    df_growth_agg_changes.reset_index().Strain.str.match("NQ393").to_list()
)

# %% [code]
df_prot = pd.read_csv(DATA / "processed_proteins.tsv", sep="\t")

# %% [code]
model = geckopy.io.read_sbml_ec_model(DATA / "eciML1515.xml.gz")

# %% [code]
model.solver = "cplex"
objective_id = "BIOMASS_Ec_iML1515_core_75p37M"
unconstrained_growth = model.slim_optimize()

# %% [code]
df_results = pd.DataFrame({"sample": [], "count": [], "average": [], "method": []})
df_results

# %% [code]
df_prot

# %% [code]
for i, (growth_mean, growth_std, sample, gdha_del) in tqdm(
    df_growth_agg_changes[
        ["growth_mean", "growth_std", "sample", "gdha_del"]
    ].iterrows(),
    total=df_growth_agg_changes[["growth_mean", "growth_std", "sample"]].shape[0],
    desc="Agg samples sensibility IIS",
):
    if len(sample) > 1:
        # group the dataframe by biological replicates (given by growth conditions)
        df_sample = (
            df_prot.loc[df_prot.Sample.isin(sample) & df_prot.in_model, ["model_id", "mmolgDW"]]
            .groupby("model_id")
            .agg([np.mean, np.std, "count"])
            .droplevel(0, axis=1)
            # use the upper limit of the CI at 0.95
            .assign(
                mmolgDW=lambda x: x["mean"] + 1.960 * x["std"] / np.sqrt(x["count"] - 1)
            )
            .reset_index()
        )
        # use the lower limit of the CI at 0.95
        growth = growth_mean - 1.960 * growth_std / sqrt(len(sample) - 1)
    else:
        df_sample = df_prot.loc[df_prot.Sample.isin(sample) & df_prot.in_model, ["model_id", "mmolgDW"]]
        growth = growth_mean
    df_sample.index = df_sample["model_id"]
    # apply concentrations (this is not inplace)
    with warnings.catch_warnings():
        warnings.simplefilter("ignore")
        model_conc = from_mmol_gDW(model, df_sample["mmolgDW"])
    # remove gdha
    if gdha_del:
        model_conc.reactions.GLUDy_REVNo1.bounds = 0, 0
        model_conc.reactions.GLUDyNo1.bounds = 0, 0
    # force a growth condition before relaxing
    model_conc.reactions.get_by_id(objective_id).lower_bound = (
        growth if growth < unconstrained_growth else unconstrained_growth
    )
    prot_candidates = [
        prot.id for prot in model_conc.proteins if prot.concentration is not None
    ]
    prot_candidates = [
        prot_id
        for prot_id in prot_candidates
        if not isnan(model_conc.proteins.get_by_id(prot_id).concentration)
    ]
    model_elastic = model_conc.copy()
    _, lp_status = get_upper_relaxation(
        model_elastic, prot_candidates, Objective_rule.MIN_ELASTIC_SUM
    )
    _ = get_upper_relaxation(model_conc, prot_candidates, Objective_rule.MIN_MILP_COUNT)
    # extract info for this sample
    if lp_status == "optimal":
        df_results = pd.concat(
            [
                df_results,
                pd.DataFrame(
                    {
                        "sample": 2 * [str(sample)],
                        "count": [
                            count_protein_saturation(model_elastic),
                            count_protein_saturation(model_conc),
                        ],
                        "average": [
                            get_average_protein_saturation(model_elastic),
                            get_average_protein_saturation(model_conc),
                        ],
                        "method": [
                            "LP first",
                            "MILP first",
                        ],
                    }
                ),
            ]
        )

# %% [code]
df_results

# %% [code]
df_results.groupby("method").mean()

# %% [code]
model.

# %% [code]
fig = px.bar(df_results, x="sample", y="average", color="method", barmode="group", template="plotly_white", height=600, width=900)

fig.update_layout(
    xaxis={"title": "Conditions"},
    yaxis={"title": "Average enzyme saturation"},
)

fig.update_xaxes(
    ticktext=ticktext,
    tickvals=tickvals,
)

fig.update_xaxes(titlefont=dict(size=21), tickfont=dict(size=16), tickangle=50)
fig.update_yaxes(titlefont=dict(size=21))

# %% [code]
fig.write_image("results_0448_avg_enzyme_saturation.pdf")

# %% [code]
px.bar(df_results, x="sample", y="count", color="method", barmode="group")

# %% [code]
model.reactions.FADRxNo1
```
