## Supplementary material for "Simultaneous application of enzyme and thermodynamic constraints to metabolic models using an updated Python implementation of GECKO": S3 File: fva_comparisons.ipynb

```
# %% [markdown]
# Figure 5 of geckopy 3.0 manuscript
Adapted from [pytfa tutorials](https://github.com/EPFL-LCSB/pytfa/blob/master/tutorials/figure_paper.py).

# %% [markdown]
##### Index
- [Flux Variability Analysis](#Flux-variability-analysis)
    - [Plain FVA](#Plain-FVA).
    - [Thermo FVA](#Thermo-FVA).
    - [Thermo concentrations FVA](#Thermo-concentrations-FVA).
    - [Proteomics FVA](#Proteomics-FVA).
    - [Thermo protein concentrations FVA](#Thermo-concentration-proteomics-FVA).
    - [Protein pool FVA](#Protein-pool-FVA).
- [Plotting](#Plotting)

# %% [code]
# import os
import errno
import logging
import re
from math import isnan, log
from os.path import join, pardir
from typing import List

import cobra
import matplotlib.pyplot as plt
import numpy as np
import pandas as pd
import pytfa
from cobra.flux_analysis.variability import flux_variability_analysis
from pytfa.analysis import variability_analysis
from pytfa.io import load_thermoDB
from pytfa.io.plotting import plot_fva_tva_comparison
from pytfa.optim.constraints import (
    BackwardDeltaGCoupling,
    BackwardDirectionCoupling,
    DisplacementCoupling,
    ForwardDeltaGCoupling,
    ForwardDirectionCoupling,
    MetaboliteConstraint,
    NegativeDeltaG,
    ReactionConstraint,
    SimultaneousUse,
)
from pytfa.optim.relaxation import relax_dgo
from pytfa.optim.utils import get_solution_value_for_variables
from pytfa.optim.variables import (
    BackwardUseVariable,
    DeltaG,
    DeltaGstd,
    ForwardUseVariable,
    LogConcentration,
)

import geckopy
from geckopy.experimental import from_copy_number
from geckopy.experimental.molecular_weights import extract_proteins
from geckopy.experimental.relaxation import (
    Objective_rule,
    apply_proteomics_elastic_relaxation,
    apply_proteomics_relaxation,
    elastic_upper_relaxation,
    get_upper_relaxation,
    relax_proteomics_greedy,
)
from geckopy.integration import relax_thermo_proteins
from geckopy.integration.pytfa import (
    adapt_gecko_to_thermo,
    get_thermo_coverage,
    get_thermo_reactions,
    translate_model_mnx_to_seed,
)

# %% [code]
CPLEX = "optlang-cplex"
GUROBI = "optlang-gurobi"
GLPK = "optlang-glpk"
ROOT = pardir
DATA = join(ROOT, "data")

# %% [markdown]
### Load the model
Load the enzyme constraint model and transform it to a thermo model (taking into account the proteins).

# %% [code]
ec_model = geckopy.io.read_sbml_ec_model(
    join(DATA, "ec_coli_core.xml"), hardcoded_rev_reactions=False
)
# ec_model = geckopy.io.read_sbml_ec_model(join(DATA, "eciML1515_seed.xml"))

# %% [code]
ec_model.solver = "cplex"

# %% [code]
ec_model.slim_optimize()
# ec_model.reactions.EX_glc__D_e.bounds = ec_model.reactions.EX_glc__D_e.flux, ec_model.reactions.EX_glc__D_e.flux

# %% [code]
## Transfrom it into thermo model
def create_thermo(ec_model):
    thermodb = load_thermoDB(join(DATA, "thermo_data.thermodb"))
    compartment_data = pytfa.io.read_compartment_data(
        join(DATA, "compartment_data.json")
    )
    #     translate_model_mnx_to_seed(
    #         ec_model, thermodb, join(DATA, "chem_xref_seedset.tsv")
    #     )
    tmodel = adapt_gecko_to_thermo(ec_model, thermodb, compartment_data, solver=CPLEX)
    return tmodel


# Info on the cobra_model
tmodel = create_thermo(ec_model)
tmodel.print_info()

# %% [markdown]
Check that the model works.

# %% [code]
tmodel.slim_optimize()

# %% [markdown]
### Flux variability analysis
Calculate variability analysis on all continuous variables

# %% [markdown]
#### Plain FVA

# %% [code]
logging.basicConfig(filename="figures_core.log", level=logging.DEBUG)

# %% [code]
ec_model.logger = logging.getLogger(__name__)
fva_fluxes = variability_analysis(ec_model.copy(), kind="reaction")

# %% [markdown]
The solver is using all available processors, so we set `n_proc` to 1 to avoid threading overhead.

# %% [markdown]
#### Thermo FVA

# %% [code]
tva_fluxes = variability_analysis(tmodel, kind="reaction")

# %% [markdown]
Save the results just in case.

# %% [code]
fva_fluxes.to_csv("fva_fluxes_core.tsv", sep="\t", index_label="reaction")
tva_fluxes.to_csv("tva_fluxes_core.tsv", sep="\t", index_label="reaction")

# %% [markdown]
#### Thermo concentrations FVA

# %% [markdown]
##### Now, the same with specific concentration data.

# %% [code]
# Add more specific concentration data
def apply_concentration_bound(met, lb, ub):
    the_conc_var = tmodel.log_concentration.get_by_id(met)
    # Do not forget the variables in the model are logs !
    the_conc_var.ub = log(ub)
    the_conc_var.lb = log(lb)

# %% [code]
tmodel = create_thermo(ec_model)

# %% [code]
apply_concentration_bound("atp_c", lb=1e-4, ub=5e-4)
apply_concentration_bound("adp_c", lb=4e-4, ub=7e-4)
apply_concentration_bound("amp_c", lb=2e-5, ub=3e-5)

tmodel.optimize()
# Perform variability analysis again
tva_fluxes_lc = variability_analysis(tmodel, kind="reactions")

# %% [code]
tva_fluxes_lc.to_csv("tva_fluxes_lc_core.tsv", sep="\t", index_label="reaction")

# %% [markdown]
#### Proteomics FVA

# %% [code]
raw_proteomics = pd.read_csv(join(DATA, "ecoli_proteomics_schmidt2016S5.tsv"))
ec_model_constrained = from_copy_number(
    ec_model.copy(),
    index=raw_proteomics["uniprot"],
    cell_copies=raw_proteomics["copies_per_cell"],
    stdev=raw_proteomics["stdev"],
    vol=2.3,
    dens=1.105e-12,
    water=0.3,
)

# %% [code]
prots_with_concentration = [
    prot.id
    for prot in ec_model_constrained.proteins
    if prot.concentration is not None and not isnan(prot.concentration)
]

# %% [code]
ec_model_constrained.reactions.BIOMASS_Ecoli_core_w_GAM.lower_bound = (
    tmodel.slim_optimize()
)
iis_lpobj = elastic_upper_relaxation(
    ec_model_constrained,
    prots_with_concentration,
    Objective_rule.MIN_ELASTIC_SUM_OBJECTIVE,
)
iis_lp = elastic_upper_relaxation(
    ec_model_constrained, prots_with_concentration, Objective_rule.MIN_ELASTIC_SUM
)
iis_milp = elastic_upper_relaxation(
    ec_model_constrained, prots_with_concentration, Objective_rule.MIN_MILP_COUNT
)

# %% [markdown]
All rules returned the same proteins.

# %% [code]
ec_model_constrained.reactions.BIOMASS_Ecoli_core_w_GAM.lower_bound = 0.0

# %% [code]
len(iis_milp)

# %% [code]
new_growth_rate, prots_relaxed = relax_proteomics_greedy(
    ec_model_constrained, minimal_growth=tmodel.slim_optimize(), protein_set=iis_milp
)

# %% [code]
tva_fluxes_prot = variability_analysis(ec_model_constrained, kind="reaction")

# %% [code]
tva_fluxes_prot.to_csv(
    "tva_fluxes_just_prot_core.tsv", sep="\t", index_label="reaction"
)

# %% [markdown]
#### Thermo concentration proteomics FVA

# %% [code]
raw_proteomics = pd.read_csv(join(DATA, "ecoli_proteomics_schmidt2016S5.tsv"))
ec_model_constrained = from_copy_number(
    ec_model.copy(),
    index=raw_proteomics["uniprot"],
    cell_copies=raw_proteomics["copies_per_cell"],
    stdev=raw_proteomics["stdev"],
    vol=2.3,
    dens=1.105e-12,
    water=0.3,
)
tmodel_prot = create_thermo(ec_model_constrained)

# %% [code]
tmodel_prot.slim_optimize()

# %% [code]
tmodel_prot.reactions.BIOMASS_Ecoli_core_w_GAM.lower_bound = tmodel.slim_optimize()
iis_milp = relax_thermo_proteins(
    tmodel_prot, prots_with_concentration, Objective_rule.MIN_MILP_COUNT
)

# %% [code]
len(iis_milp[0])

# %% [code]
raw_proteomics = pd.read_csv(join(DATA, "ecoli_proteomics_schmidt2016S5.tsv"))
ec_model_constrained = from_copy_number(
    ec_model.copy(),
    index=raw_proteomics["uniprot"],
    cell_copies=raw_proteomics["copies_per_cell"],
    stdev=raw_proteomics["stdev"],
    vol=2.3,
    dens=1.105e-12,
    water=0.3,
)
tmodel_prot = create_thermo(ec_model_constrained)
for prot in iis_milp[0]:
    tmodel_prot.proteins.get_by_id(prot).concentration = None

# %% [code]
tmodel_prot.optimize()

# %% [code]
# Perform variability analysis again
tva_fluxes_prot = variability_analysis(tmodel_prot, kind="reactions")

# %% [code]
tva_fluxes_prot.to_csv("tva_fluxes_prot_core.tsv", sep="\t", index_label="reaction")

# %% [markdown]
#### Protein pool FVA

# %% [code]
pool_model = geckopy.io.read_sbml_ec_model(
    join(DATA, "ec_coli_core.xml"), hardcoded_rev_reactions=False
)
pool_model.solver = "cplex"
pool_model.reactions.EX_glc__D_e.bounds = (
    ec_model.reactions.EX_glc__D_e.flux,
    ec_model.reactions.EX_glc__D_e.flux,
)
pool_model.slim_optimize()

# %% [code]
df = extract_proteins(pool_model)

# %% [code]
for row in df.itertuples():
    pool_model.proteins.get_by_id(row[2]).mw = row[3]

# %% [code]
pool_model.constrain_pool(
    0.448,
    0.6,
    1.0,
)

# %% [code]
pool_model.slim_optimize()

# %% [markdown]
We take the smaller set.

# %% [code]
pool_model.reactions.BIOMASS_Ecoli_core_w_GAM.lower_bound = 0

# %% [code]
pool_model.slim_optimize()

# %% [code]
pool_model.logger = logging.getLogger(__name__)
tva_fluxes_pool = variability_analysis(pool_model)

# %% [code]
tva_fluxes_pool.to_csv("tva_fluxes_pool_core.tsv", sep="\t", index_label="reaction")

# %% [markdown]
### Plotting

# %% [code]
def plot_stacked_negative(
    dfs=List[pd.DataFrame],
    labels=List[str],
    colors=List[str],
    widths=List[float],
    width_col=0.12,
):
    """Plot stacked negative barplot."""
    n_rows = dfs[0].shape[0]
    ind = np.arange(n_rows)
    fig, ax = plt.subplots(figsize=(18, 6))

    for df, label, color, width in zip(dfs, labels, colors, widths):
        p1 = ax.bar(ind + width, df.maximum, width_col, label=label, color=color)
        p2 = ax.bar(ind + width, df.minimum, width_col, color=color)

    ax.axhline(0, color="#221c22", linewidth=0.8, linestyle="--")
    ax.set_ylabel("Fluxes")
    ax.set_xticks(ind)
    ax.set_xticklabels(dfs[0].index, rotation=-40)
    ax.legend()


def apply_rev(reac_id, df, model):
    direction = "maximum"
    rev_id = re.sub(REV_PATTERN, r"_REV\1", reac_id)
    try:
        rev_id = model.reactions.get_by_id(rev_id).id
    except KeyError:
        rev_id = reac_id
        direction = "minimum"
    return -df.loc[rev_id, direction]


def apply_df_rev(df, model):
    df["minimum"] = [apply_rev(reac_id, df, ec_model) for reac_id in df.index]

# %% [markdown]
Load previously generated output.

# %% [code]
fva_fluxes = pd.read_csv("fva_fluxes_core.tsv", sep="\t", index_col="reaction").assign(
    method="FBA"
)
tva_fluxes = pd.read_csv("tva_fluxes_core.tsv", sep="\t", index_col="reaction").assign(
    method="Thermo"
)
tva_fluxes_lc = pd.read_csv(
    "tva_fluxes_lc_core.tsv", sep="\t", index_col="reaction"
).assign(method="Thermo + Metabolomics")
tva_fluxes_prot = pd.read_csv(
    "tva_fluxes_prot_core.tsv", sep="\t", index_col="reaction"
).assign(method="Thermo + Proteomics")
tva_fluxes_just_prot = pd.read_csv(
    "tva_fluxes_just_prot_core.tsv", sep="\t", index_col="reaction"
).assign(method="Proteomics")
tva_fluxes_pool = pd.read_csv(
    "tva_fluxes_pool_core.tsv", sep="\t", index_col="reaction"
).assign(method="Pool constraint")

# %% [code]
dfs = (
    fva_fluxes,
    tva_fluxes,
    tva_fluxes_lc,
    tva_fluxes_prot,
    tva_fluxes_pool,
    tva_fluxes_just_prot,
)

# %% [code]
dfsub = pd.concat(
    [
        fva_fluxes,
        tva_fluxes.rename({"minimum": "min_prot", "maximum": "max_prot"}, axis=1),
    ],
    axis=1,
)
reacs_to_plot = (
    dfsub.apply(lambda s: s.maximum - s.minimum + s.min_prot - s.max_prot, axis=1)
    .sort_values(ascending=False)[:20]
    .index
)

# %% [code]
dfsub = pd.concat(
    [
        fva_fluxes,
        tva_fluxes_prot.rename({"minimum": "min_prot", "maximum": "max_prot"}, axis=1),
    ],
    axis=1,
)
reacs_to_plot.append(
    dfsub.apply(lambda s: s.minimum - s.min_prot, axis=1).sort_values()[:10].index
)

# %% [code]
reacs_to_plot = [
    "ATPS4r",
    "NADTRHD",
    "PPCK",
    "ADK1",
    "FBA",
    "G6PDH2r",
    "PFK",
    "PPS",
    "PYK",
    "TPI",
]

# %% [markdown]
Figure 5 BEC of [Sánchez et al., 2017](https://www.embopress.org/doi/full/10.15252/msb.20167411).

# %% [code]
df_plot = pd.concat(
    [
        abs(df.maximum - df.minimum)
        .apply(lambda x: x if x < 2000 else 2000)
        .to_frame()
        .assign(method=df.iloc[0, 2])
        .rename({0: "Flux"}, axis=1)
        for df in dfs
    ]
)

# %% [code]
df_plot

# %% [code]
abs(tva_fluxes_pool.maximum - tva_fluxes_pool.minimum).apply(
    lambda x: x if x < 2000 else 2000
).max()

# %% [code]
import plotly.express as px

# %% [code]
fig = px.histogram(
    df_plot[df_plot.Flux.abs() > 1e-7],
    x="Flux",
    color="method",
    cumulative=True,
    nbins=1000,
    marginal="box",
    barmode="overlay",
    opacity=0.1,
    width=800,
    height=600,
)
fig.update_layout(
    legend=dict(
        x=0.7,
        y=0.05,
        traceorder="normal",
        font=dict(family="sans-serif", size=12, color="black"),
    )
)

# %% [code]
fig = px.box(
    df_plot[df_plot.Flux.abs() > 1e-7],
    x="Flux",
    y="method",
    color="method",
    log_x=True,
    template="plotly_white",
    width=800,
    height=600,
)
fig.update_layout(showlegend=False, yaxis={"tickangle": -45})

fig.update_xaxes(title_font={"size": 20}, tickfont=dict(size=14))
fig.update_yaxes(tickfont=dict(size=16), title=None)
fig

# %% [code]
fig.write_image("/home/georg/Downloads/fva_comp.pdf")

# %% [code]
df_plot_b = pd.concat(dfs)
df_plot_b["Reaction"] = df_plot.index
# df_plot_b.loc[(df_plot_b.minimum - df_plot_b.maximum).abs() < 1e-4, "minimum"] = 0.
df_plot_b = df_plot_b.melt(["method", "Reaction"], var_name="Sense", value_name="Flux")
df_plot_b

# %% [code]
fig = px.bar(
    df_plot_b[df_plot_b.Reaction.isin(reacs_to_plot)],
    x="Reaction",
    y="Flux",
    color="method",
    labels="Sense",
    barmode="group",
    template="plotly_white",
    width=800,
    height=600,
)
fig.update_layout(
    legend=dict(
        x=0.03,
        y=1.0,
        traceorder="normal",
        font=dict(family="sans-serif", size=16, color="black"),
    )
)
fig.update_xaxes(title_font={"size": 20}, tickfont=dict(size=16))
fig.update_yaxes(title_font={"size": 20}, tickfont=dict(size=14))
fig

# %% [code]
fig.write_image("/home/georg/Downloads/fva_comp_reacs.pdf")

# %% [code]
[reac.id for reac in ec_model.reactions if reac.lower_bound == 0]

# %% [code]
47 / 95 * 100
```
